# fMRI reveals language-specific predictive coding during naturalistic sentence comprehension

**DOI:** 10.1101/717512

**Authors:** Cory Shain, Idan Asher Blank, Marten van Schijndel, William Schuler, Evelina Fedorenko

## Abstract

Much research in cognitive neuroscience supports prediction as a canonical computation of cognition across domains. Is such predictive coding implemented by feedback from higher-order domain-general circuits, or is it locally implemented in domain-specific circuits? What information sources are used to generate these predictions? This study addresses these two questions in the context of language processing. We present fMRI evidence from a naturalistic comprehension paradigm (1) that predictive coding in the brain’s response to language is domain-specific, and (2) that these predictions are sensitive both to local word co-occurrence patterns and to hierarchical structure. Using a recently developed continuous-time deconvolutional regression technique that supports data-driven hemodynamic response function discovery from continuous BOLD signal fluctuations in response to naturalistic stimuli, we found effects of prediction measures in the language network but not in the domain-general multiple-demand network, which supports executive control processes and has been previously implicated in language comprehension. Moreover, within the language network, surface-level and structural prediction effects were separable. The predictability effects in the language network were substantial, with the model capturing over 37% of explainable variance on held-out data. These findings indicate that human sentence processing mechanisms generate predictions about upcoming words using cognitive processes that are sensitive to hierarchical structure and specialized for language processing, rather than via feedback from high-level executive control mechanisms.

## Introduction

The human brain is an efficient prediction engine (James, 1890). Facilitation in processing expected information, as well as processing costs of violated expectations, have been reported in many domains. In the domain of language comprehension, various results show that listeners and readers actively predict upcoming words and structures (e.g., Kutas & Hillyard, 1984; MacDonald et al., 1994; Tanenhaus et al., 1995; Rayner et al., 2004; Frank & Bod, 2011; Smith & Levy, 2011, 2013; Staub & Benetar, 2013; Frank et al., 2015; Kuperberg & Jaeger, 2016). However, the cognitive and neural mechanisms that support predictive language processing are not well understood. Under one widely held view, predictive language processing is implemented by domain-general executive (inhibitory control and working memory) resources. This perspective receives support from numerous studies showing that prediction effects during language comprehension are absent or less pronounced for populations with reduced executive resources, such as children, older individuals, and non-native speakers (e.g., Federmeier et al., 2002; Federmeier & Kutas, 2005; Dagerman, et al., 2006; Federmeier et al., 2010; Mani & Huettig, 2012; Wlotko & Federmeier, 2012; Martin et al., 2013; Kaan, 2014; Mitsugi & Macwhinney, 2016; Gambi et al., 2018; Payne & Federmeier, 2018; cf. Dave et al., 2018; Havron et al., 2019). Furthermore, several neuroimaging studies have reported sensitivity to linguistic manipulations in what appear to be cortical regions thought to support domain-general executive function (e.g., Kaan & Swaab; 2002; Kuperberg et al., 2003; Novick et al., 2005; Rodd et al., 2005; Novais-Santos, 2007; January et al., 2009; Peelle et al., 2010; Rogalsky & Hickok, 2011; Nieuwland et al., 2012; Wild et al., 2012; McMillan et al., 2012, 2013), suggesting that such regions may also be implicated in language processing, including perhaps prediction. These results have led some to conclude that predictive coding for language is implemented by domain-general executive control resources (Linck et al., 2014; Huettig & Mani, 2016; Pickering & Gambi, 2018; Strijkers et al., 2019).

However, this interpretation is subject to several objections. First, most prior work on linguistic prediction has relied on behavioral and electrophysiological measures which are well suited for identifying global response patterns but cannot spatially localize the source of these effects in the brain to a certain functional region or network (e.g., Mather et al., 2013). Second, the (alleged) between-population differences in prediction noted above are consistent with accounts that do not directly invoke executive resources, including (1) possible qualitative differences between populations in the kind of information that is being predicted and the consequent need for population-specific norms to detect prediction effects, or (2) differences in how often predictions are correct, which may modulate the likelihood of engaging in predictive behavior (see Ryskin et al., this issue, for discussion). And third, past studies that did employ neuroimaging tools with high spatial resolution and consequently reported linguistic prediction responses

– typically neural response increases for violations of linguistic structure – localized to executive control regions (e.g., Newman et al., 2001; Kuperberg et al., 2003; Nieuwland et al., 2012; Schuster et al., 2016) may have been influenced by task artifacts; indeed, some have argued that artificially constructed laboratory stimuli and tasks increase general cognitive load in comparison to naturalistic language comprehension (e.g., Blanco-Elorietta & Pylkkänen, 2017; Blank & Fedorenko, 2017; Campbell & Tyler, 2018; Wehbe et al., submitted; Diachek et al., 2019). To ensure that findings from the laboratory paradigms truly reflect the cognitive phenomenon of interest, it is important to validate them in more naturalistic experimental settings that better approximate the typical conditions of human sentence comprehension (Hasson & Honey, 2012; Hasson et al., 2018).

Despite the growing number of fMRI studies of naturalistic language comprehension (e.g., Speer et al., 2007; Yarkoni et al., 2008; Speer et al., 2009; Whitney et al., 2009; Wehbe et al., 2014; Hale et al., 2015; Henderson et al., 2015, 2016; Huth et al., 2016; Sood & Sereno, 2016; Brennan, 2016; Desai et al., 2016; de Heer et al., 2017, Dehghani et al., 2017; Bhattasali et al., 2018), only a handful have directly investigated effects of word predictability (Willems et al., 2015; Brennan et al., 2016; Henderson et al.,

2016; Lopopolo et al., 2017; see **Table 1** for summary), a well-established predictor of behavioral measures in naturalistic language comprehension (Demberg & Keller, 2008; Frank & Bod, 2011; Smith & Levy, 2013; van Schijndel & Schuler, 2015). These previous naturalistic studies of linguistic prediction effects in the brain – using estimates of prediction effort such as *surprisal* (Hale, 2001; Levy, 2008), the negative log probability of a word given its context, or *entropy* (Hale, 2006), an information-theoretic measure of the degree of constraint placed by the context on upcoming words – have yielded mixed results on the existence, type, and functional location of such effects. For example, of the lexicalized and unlexicalized (part-of-speech) bigram and trigram models of word surprisal explored in Brennan et al. (2016), only part-of-speech bigrams positively modulated neural responses in most regions of the functionally localized language network. Lexicalized bi- and trigrams and part-of-speech trigrams yielded generally null or negative results (16 out of 18 comparisons). By contrast, Willems et al. (2015) found lexicalized trigram effects in regions typically associated with language processing (e.g., anterior and posterior temporal lobe). In addition, Willems et al. (2015) and Lopopolo et al. (2017) found prediction effects in regions that are unlikely to be specialized for language processing, including (aggregating across both studies) the brain stem, amygdala, putamen, and hippocampus, as well as in superior frontal areas more typically associated with domain-general executive functions like self-awareness and coordination of the sensory system (Goldberg et al., 2006). It is therefore not yet clear whether predictive coding for language relies on domain-general mechanisms in addition to, or instead of, language-specific ones, especially in naturalistic contexts.

In addition to questions about the functional localization of linguistic prediction, substantial prior work has also investigated the structure of the predictive model, seeking to shed light on the nature of linguistic representations in the mind. If effects from theoretical constructs like hierarchical natural language syntax can be detected in online processing measures, this would constitute evidence that such constructs are present in human mental representations and used to comprehend language. This position is widely supported by behavioral and electrophysiological experiments using constructed stimuli (see Lewis & Phillips, 2015 for review) and by some behavioral (Roark et al., 2009; Fossum & Levy, 2012; van Schijndel & Schuler, 2015; Shain et al., 2016), electrophysiological (Brennan & Hale, 2019) and neuroimaging (Brennan et al., 2016) experiments using naturalistic stimuli. However, other naturalistic studies reported null or negative syntactic effects (Frank & Bod, 2011; van Schijndel & Schuler, 2013; Shain & Schuler, 2018 *contra* Shain et al., 2016), or mixed syntactic results within the same set of experiments (Demberg & Keller, 2008; Henderson et al, 2016), leading some to argue that the representations used for language comprehension (in the absence of task artifacts from constructed stimuli) contain little hierarchical structure (Frank & Christiansen, 2018). Furthermore, the few naturalistic fMRI studies that have explored structural prediction effects have yielded inconsistent localization of these effects. For example, Brennan et al. (2016) found context-free grammar surprisal effects throughout the functional language network except in inferior frontal gyrus, whereas inferior frontal gyrus is the only region in which Henderson et al. (2016) found such effects.

The current study used fMRI to determine whether a signature of predictive coding during language comprehension – increased response to less predictable words, i.e. surprisal (e.g., Smith & Levy, 2013) – is primarily evident during naturalistic sentence processing in (1) the domain-specific, fronto-temporal language (LANG) network (Fedorenko et al., 2011), or (2) the domain-general, fronto-parietal multiple demand (MD) network (Duncan, 2010). The MD network supports executive functions (e.g., inhibitory control, attentional selection, conflict resolution, maintenance and manipulation of task sets) across both linguistic and non-linguistic tasks (e.g., Duncan & Owen, 2000; Fedorenko et al., 2013; Hughdahl et al., 2015; for discussion, see: Fedorenko, 2014) and has been shown to be sensitive to surprising events (Corbetta & Shulman, 2002).

On the one hand, given that the language network plausibly stores linguistic knowledge, including the statistics of language input, it might directly carry out predictive processing. Such a result would align with a growing body of cognitive neuroscience research supporting prediction as a “canonical computation” (Keller & Mrsic-Flogel, 2018) locally implemented in domain-specific circuits (Montague et al, 1996; Rao & Ballard, 1999; Alink et al., 2010; Bubic et al., 2010; Bastos et al., 2012; Wacogne et al., 2011, 2012; Singer et al., 2018). This hypothesis is also supported by prior findings of linguistic prediction effects in portions of the language network (Bonhage et al., 2015; Willems et al., 2015; Brennan et al., 2016; Henderson et al., 2016; Lopopolo et al., 2017; Matchin et al., 2018).

On the other hand, given that the MD network has been argued to encode predictive signals across domains and relay them as feedback to other regions (Strange et al., 2005; Cristescu et al., 2006; Egner et al., 2008; Wacogne et al., 2011; Chao et al., 2018), it might be recruited to predict upcoming words and structures in language. There is an extensive literature on neural signatures of prediction, such as activity associated with prediction errors, in brain regions that appear to belong to the MD network, including bilateral areas in the dorsolateral pre-frontal cortex, the inferior frontal gyrus, the anterior cingulate cortex, and the parietal lobe (for a review, see Dehaene et al., 2015; for a meta-analysis, see D’Astolfo & Rief, 2017). These areas are sensitive to rule violations in non-linguistic sequences, including hierarchically structured ones, in different sensory domains (e.g., auditory and visual; Bekinschtein et al., 2009; Ahlheim et al., 2014; Uhrug et al., 2014; Wang et al., 2015; Wang et al., 2017; Chao et al., 2018). In addition, they are recruited during learning of structured sequences in the motor domain (Bischoff-Grethe et al., 2004; Eickhoff et al., 2010). Beyond representing deterministic rules, such regions are also engaged in probabilistic predictions (Strange et al., 2005; Meyniel & Dehaene, 2017). Such predictions can be based on either inferring a generative model underlying the input sequence (Gläscher et al., 2010; Schapiro et al., 2013) or on reward contingencies (Koch et al., 2008; Zarr & Brown, 2016; Alexander & Brown, 2018; for a review, see: Rushworth & Behrens, 2008).

There are two main hypotheses in the contemporary literature that link predictive processing in the MD network with increased activity to more surprising words. First, the MD network might provide additional resources (“cognitive juice”) to various cognitive processes, including language. Under this scenario, MD regions might “come to the rescue” of the language network when processing demands are increased, which would be the case when surprisal is higher. Indeed, prior work suggests that the MD network could be recruited when language processing becomes effortful, e.g., under acoustic (Adank, 2012; Hervais-Adelman et al., 2012; Wild et al., 2012; Scott & McGettigan, 2013; Vaden et al., 2013) or syntactic (Kuperberg et al., 2003; Nieuwland et al., 2013) noise; in healthy aging (for reviews, see Wingfield & Grossman, 2006; Shafto & Tyler, 2014); during recovery from aphasia (Brownsett et al., 2014; Geranmayeh et al., 2014, 2016, 2017; Meier et al., 2016; Sims et al., 2016; Hartwigsen, 2018); and in L2 processing and multi-lingual control (e.g., Wartenburger et al., 2003; Rüschemeyer et al., 2005; Yokoyama et al., 2006; de Bruin et al., 2014; Grant, Fang, & Li, 2015; Kim et al., 2016; for reviews, see Perani & Abutalebi, 2005; Sakai, 2005; Abutalebi, 2008; Kotz, 2009; Hervais-Adelman, Moser-Mercer, Golestani, 2011; Pliatsikas & Luk, 2016). Second, the MD network, especially in the prefrontal cortex, may construct abstract representations of context, which serve as working memory for guiding behavior (Alexander & Brown, 2018). The main goal of such representations is to minimize prediction errors in other brain regions, so these representations are communicated in a top-down manner to the language network or other domain-specific networks (e.g., sensory areas). Such high-level, abstract predictive signals are potentially useful because they could perhaps “explain away” some more local prediction errors computed in the language network (e.g., in a sentence like “the cat that the dog chased on the balcony escaped”, the verb “escaped” might be unexpected based on the local context of the previous few words, but its occurrence could be explained away by a more global and abstract representation that looks farther into the past and predicts a verb for “the cat” in the main clause). In essence, then, signals from the MD network could bias representations in the language network in favor of the features that are most relevant in a given context (for a similar reasoning for sensory cortices, see Miller & Cohen, 2001; Sreenivasan et al., 2014; D’Esposito & Postle, 2015). However, these higher-level predictions are still sometimes incorrect, and when errors propagate back to the MD network, its regions would be triggered to adjust their predictive model in order to minimize future errors. This “model revision” process may register as increased neural processing (Chao et al., 2018).

Prior fMRI studies using hand-constructed sentences to probe effects of linguistic expectation have not yielded a clear answer as to the mechanisms – language-specific vs. domain-general – that support linguistic prediction. Numerous such studies have observed responses in areas of the language network to manipulations of word predictability (Kuperberg et al., 2000; Baumgaertner et al., 2002; Kiehl et al., 2002; Friederici et al., 2003; Gold et al., 2006; Obleser et al., 2007; Dien et al., 2008; Obleser et al., 2009; Bonhage et al., 2015; Schuster et al., 2016; Hartwigsen et al., 2017; Matchin et al., 2018; Schuster et al., 2019). However, many studies have also reported linguistic prediction effects in frontal, parietal, and cingulate cortical regions typically associated with the MD network (Kuperberg et al., 2000; Baumgaertner et al., 2002; Gold et al., 2006; Bonhage et al., 2015; Hartwigsen et al., 2017), as well as in other parts of the brain like the fusiform gyrus (Kuperberg et al., 2000; Gold et al., 2006) and the cerebellum (Lesage et al., 2017). Although it is certainly possible that predictive coding for language is carried out by both the LANG and the MD networks, with additional contributions from other brain areas, it is important to ensure that the foregoing results are not due to task artifacts induced by the use of artificially constructed stimuli (see Discussion), through validation of these findings in more naturalistic comprehension conditions (Hasson et al., 2018).

To distinguish the hypotheses above in a naturalistic comprehension paradigm, we searched for neural responses in LANG vs. MD regions to the contextual predictability of words as estimated by two model implementations of surprisal: a surface-level 5-gram model and a hierarchical probabilistic context-free grammar (PCFG) model. *N*-gram surprisal estimates are sensitive to word co-occurrence patterns but are limited in their ability to model hierarchical natural language syntax, since they contain no explicit representation of grammatical categories or syntactic composition and have limited memory for preceding words in the sentence (in our case, up to four preceding words). PCFG surprisal estimates, by contrast, are based on structured syntactic representations of the unfolding sentence but do not directly encode surface-level word co-occurrence patterns. Correlations between each of these measures and human neural responses would shed light on the relative importance assigned to these two information sources (word co-occurrences and syntactic structures) in computing predictions about upcoming words. Although surprisal is not the only extant measure of linguistic prediction (others include PCFG entropy, Roark et al., 2009; entropy reduction, Hale, 2006; and successor surprisal, Kliegl et al., 2006), surprisal has received extensive consideration in the experimental literature (e.g., Demberg and Keller, 2008; Frank & Bod, 2011; Fossum & Levy, 2012; Frank et al, 2015; van Schijndel and Schuler, 2015; Brennan et al., 2016; Henderson et al., 2016; Brennan & Hale, 2019; Shain, 2019). We did not consider these related measures in order to avoid excessive statistical comparisons.

Note that by estimating prediction effects using surprisal, we are implicitly assuming a notion of linguistic prediction as a distributed *pre-activation* process, following e.g. Kuperberg & Jaeger (2016), rather than as an all-or-nothing commitment to a specific upcoming word. Thus, we are investigating the degree to which the statistics of the local lexical (*n*-gram) and structural (PCFG) linguistic context modulate the sentence processing response, and where in the brain this modulation occurs. We leave aside questions about the underlying mechanisms by which these effects arise: e.g. the extent to which they are “active” or “passive”, or the extent to which integrative structure-building operations (e.g. composing words into syntactic constituents, constructing dependencies, etc.) underlie the observed facilitation effects (Altmann, 1998; Hale, 2014). See Discussion for elaboration on this point.

To avoid the problem of reverse inference from anatomy to function (Poldrack 2006, 2011; **Figure 1**), we functionally defined the LANG and MD networks in each individual participant using an independent “localizer” task (Saxe et al., 2006; Fedorenko et al., 2010), and then examined the response of those functional regions to each estimate of surprisal. Our results show significant independent effects of 5-gram and PCFG surprisal in LANG, but no such effects in MD, as well as significant differences in surprisal effect sizes between the two networks. This finding supports the hypothesis that predictive coding for language is primarily carried out by language-specialized rather than domain-general cortical circuits and exploits both surface-level and structural cues.

## Materials and Methods

### General Approach

Several features set the current study apart from prior cognitive neuroscience investigations of linguistic prediction in naturalistic stimuli.

First, we used naturalistic language stimuli rather than controlled stimuli constructed for a particular experimental goal. Naturalistic stimuli improve ecological validity compared to isolated constructed stimuli, which may introduce task artifacts that do not generalize to everyday cognition (Demberg & Keller, 2008; Hasson & Honey, 2012; Richlan et al., 2013; Schuster et al., 2016; Campbell & Tyler, 2018), and prior work indicates that naturalistic stimuli yield more reliable BOLD signals than artificial tasks (Hasson et al., 2010). Minimizing such artifacts is crucial in studies of the MD network, which is highly sensitive to task variables (Miller & Cohen, 2001; Sreenivasan et al., 2014; D’Esposito & Postle, 2015; Diachek et al., 2019).

Second, we used participant-specific functional localization to identify regions of interest constituting the LANG and MD networks (Fedorenko et al., 2010). This approach is crucial because many functional regions do not exhibit a consistent mapping onto macro-anatomical landmarks (Frost & Goebel, 2012), especially in the frontal (Amunts et al., 1999; Tomaiuolo et al., 1999), temporal (Jones and Powell, 1970; Gloor, 1997; Wise et al., 2001) and parietal (Caspers et al., 2006; Caspers et al., 2008; Scheperjans et al., 2008) lobes, which house the language and MD networks. Due to this inconsistent functional-to-anatomical mapping, a given stereotactic coordinate might belong to the language network in some participants but to the MD network in others, as is indeed the case in our sample (**Figure 1**) (see also Fedorenko et al., 2012a; Blank et al., 2017; Fedorenko & Blank, submitted). Such inter-individual variability severely compromises the validity of both anatomical localization (Juch et al., 2005; Poldrack, 2006; Fischl et al., 2007; Frost and Goebel, 2012; Tahmasebi et al., 2012) and group-based functional localization (Saxe et al., 2006; Fedorenko and Kanwisher, 2009): these approaches risk both decreased sensitivity (i.e. failing to identify a functional region due to insufficient spatial alignment across participants) and decreased functional resolution (i.e. mistaking two functionally distinct regions for a single region due to apparent spatial overlap across the sample). In contrast, participant-specific functional localization allows us to pool data from a given functional region across participants even in the absence of perfect anatomical alignment and is therefore better suited for the kind of questions we study here (Nieto-Castañón & Fedorenko, 2012). Both networks we probe here have been extensively functionally characterized in prior work, so responses to linguistic surprisal therein can be taken to index the engagement of linguistic processing mechanisms vs. domain-general executive mechanisms (e.g., Mather et al., 2013).

**Figure 1:**
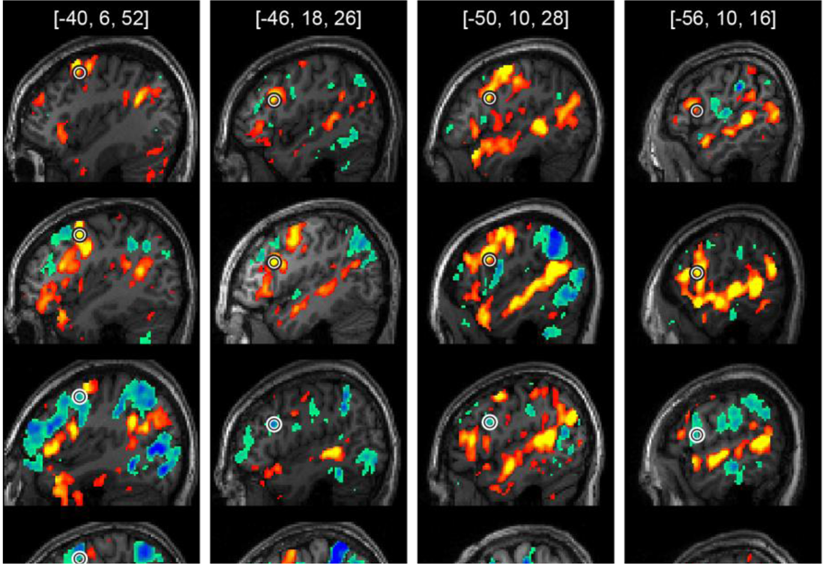
Inter-individual variability in the mapping of function onto anatomy. Each column demonstrates variability in a different coordinate in MNI space, specified at the top (in mm). For each coordinate, sagittal T1 slices from four participants are shown, with the coordinate circled on each slice (participants differ across columns). In each case, the top two participants show a Sentences > Nonwords effect in this coordinate (colored in red-yellow), whereas the bottom two participants show the opposite, Nonwords > Sentences effect in this same coordinate (colored in green-blue). In all cases, the effect size of the circled coordinate is strong enough to be included among the participant-specific fROIs. Other voxels exhibiting strong contrast effects in the localizer task (namely, among the top 10% of voxels across the neocortical gray matter) are superimposed onto the anatomical slices, in color. Colorbars show p-values associated with each of the two localizer contrasts.

Third, we analyzed the BOLD times-series using a recently developed statistical framework – continuous-time deconvolutional regression (CDR; Shain & Schuler, 2018, 2019) – that is designed to overcome problems in hemodynamic response modeling that are presented by naturalistic experiments. The variable spacing of words in naturalistic language prevents direct application of discrete-time, data-driven techniques for hemodynamic response function (HRF) discovery, such as finite impulse response modeling (FIR) or vector autoregression. Because CDR is a parametric continuous-time deconvolutional method, it can infer the hemodynamic response directly from naturalistic time series, without distortionary preprocessing steps such as predictor interpolation (cf. Huth et al., 2016). Thus, unlike prior naturalistic fMRI studies of prediction effects in language processing (**Table 1**), we do not assume the shape of the HRF.

Fourth, unlike studies in **Table 1**, we evaluated hypotheses using non-parametric statistical tests of model fit to held-out (out-of-sample) data, an approach which builds external validity directly into the statistical test and should thereby improve replicability (e.g., Demšar, 2006).

**Table 1.** Previous fMRI studies of prediction effects in naturalistic sentence comprehension.

| Study | # Participants | Stimulus length | HRF model | Functional localization | Out-of-sample evaluation |
| --- | --- | --- | --- | --- | --- |
| Willems et al., 2015 | 24 | 19 min | Canonical | No | No |
| Brennan et al., 2016 | 26 | 12 min | Canonical | Yes | No |
| Henderson et al., 2016 | 40 | 22 paragraphs | Canonical | No | No |
| Lopopolo et al., 2017 | 22 | 19 min | Canonical | No | No |
| **Current study** | **78** | **13.5 min (avg per participant)** | **Data-driven (CDR)** | **Yes** | **Yes** |

Finally, to our knowledge, this is the largest fMRI investigation to date (78 subjects) of prediction effects in naturalistic language comprehension.

## Experimental Design

### Participants

Seventy-eight native English speakers (30 males), aged 18-60 (*M*±*SD* = 25.8±9, *Med*±*SIQR* = 23±3), from MIT and the surrounding Boston community participated for payment. Each participant completed a passive story comprehension task (the critical experiment) and a functional localizer task designed to identify the language and MD networks.

Sixty-nine participants (88%) were right-handed, as determined by either the Edinburgh handedness inventory (*n*=66) (Oldfield, 1971) or self-report (*n*=11) (handedness data were not collected for one participant). Eight participants were left-handed, but seven of these showed typical left-lateralized language activations, as determined by examining their activation patterns for the language localizer task (see below); the remaining participant had a right-lateralized language network. We chose to include the latter participant’s data in the analyses, to err on the conservative side, and to be able to generalize the results to the population at large (see Willems et al., 2014, for discussion).

All participants gave informed consent in accordance with the requirements of MIT’s Committee on the Use of Humans as Experimental Subjects (COUHES).

### Stimuli and procedure

The localizer task and critical (story comprehension) experiment were run either in the same scanning session (67 participants) or in two separate sessions (11 participants, who have performed the localizer task while participating in other studies; see Mahowald & Fedorenko, 2016, for evidence of high stability of language localizer activations across sessions). For the critical experiment, each participant listened to one or more stories (one story: *n*=34; two stories: *n*=14; three stories: *n*=13; four stories: *n*=2; five stories: *n*=4; six stories: *n*=5; seven stories: *n*=1; or eight stories: *n=5*). In each session, participants performed a few other, unrelated tasks, with scanning sessions lasting approximately 2h.

*Localizer task.* We used a single localizer task to identify functional regions of interest in both the language and MD networks, using opposite task contrasts across these networks as described below. This task, which has been described in more detail elsewhere (Fedorenko et al., 2010), consisted of reading sentences and lists of unconnected, pronounceable nonwords in a standard two-condition blocked design with a counterbalanced order across runs. Stimuli were presented one word / nonword at a time. The majority of participants (*n*=60) read these materials passively (and pressed a button at the end of each trial, to sustain alertness); for some participants (*n*=18), every trial ended with a memory probe item, and they had to indicate via a button press whether or not this probe had appeared in the preceding sentence / nonwords sequence. In addition, different participants performed versions of the task differing slightly in stimulus timing, number of blocks, etc., i.e. features that that do not affect the robustness of the contrast (e.g., Fedorenko et al., 2010; Mahowald & Fedorenko, 2016) (for experimental parameters, see Table 2). A version of this localizer is available at https://evlab.mit.edu/funcloc/download-paradigms.

**Table 2.** Experimental parameters for the different versions of the localizer task. *Used for the purposes of another experiment; see (Fedorenko et al., 2010).

|  | Version |  |  |  |
| --- | --- | --- | --- | --- |
|  | A | B | C | D |
| Number of participants | 60 | 6 | 5 | 7 |
| Task (Passive Reading / Memory) | PR | M | M | M |
| Words / nonwords per trial | 12 | 12 | 8 | 12 |
| Trial duration (ms) | 6,000 | 6000 | 4,800 | 6000 |
| Fixation | 100 | 300 | 300 | 300 |
| Presentation of each word / nonword | 450 | 350 | 350 | 350 |
| Probe (M) + button press (M/PR) | 400 | 1000 | 1350 | 1000 |
| Fixation | 100 | 500 | 350 | 500 |
| Trials per block | 3 | 3 | 5 | 3 |
| Block duration (s) | 18 | 18 | 24 | 18 |
| Blocks per condition (per run) | 8 | 8 | 4 | 6 |
| Conditions | Sentences<br>Nonwords | Sentences<br>Nonwords | Sentences<br>Nonwords<br>Word-lists* | Sentences<br>Nonwords<br>Word-lists* |
| Fixation block duration (s) | 14 | 18 | 16 | 18 |
| Number of fixation blocks | 5 | 5 | 3 | 4 |
| Total run time (s) | 358 | 378 | 336 | 396 |
| Number of runs | 2 | 2 | 3-4 | 2-3 |

To identify language regions, we used the contrast *sentences > nonwords*. This contrast targets higher-level aspects of language, to the exclusion of perceptual (speech / reading) and motor-articulatory processes (for discussion, see Fedorenko & Thompson-Schill, 2014; or Fedorenko, in press). Critically, this localizer has been extensively validated over the past decade across diverse parameters: it generalizes across task (passive reading vs. memory probe), presentation modality (visual vs. auditory), and materials (e.g., Fedorenko et al., 2010; Braze et al., 2011; Vagharchakian et al., 2012), including both coarser contrasts (e.g., between natural speech and an acoustically degraded control: Scott et al., 2017) and narrower contrasts (e.g., between lists of unconnected, real words and nonwords lists, or between sentences and lists of real words: Fedorenko et al., 2010; Blank et al., 2016). Whereas there are many potential differences (linguistic and otherwise) between the processing of sentences vs. nonwords, all regions localized with the *sentences > nonwords* contrast show a similar response profile: on the one hand, they exhibit sensitivity to various aspects of linguistic processing, including (but not limited to) lexical, phrasal, and sentence-level semantic and syntactic processing (e.g., Fedorenko et al., 2012b, 2018; Blank et al., 2016; Mollica et al., 2018; Blank & Fedorenko, submitted; similar patterns obtain in electrocorticographic data with high temporal resolution: Fedorenko et al., 2016). On the other hand, they show robust language-selectivity in their responses, with little or no response to non-linguistic tasks, including domain-general contrasts targeting, e.g., working memory or inhibitory control (Fedorenko et al., 2011, 2012a). In other words, the localizer shows both convergent construct validity with other linguistic contrasts and discriminant construct validity against non-linguistic contrasts. Moreover, the functional network identified by this contrast is internally synchronized yet strongly dissociated from other brain networks during naturalistic cognition (e.g., Blank et al, 2014; Paunov et al., 2019; for evidence from inter-individual effect-size differences, see: Mineroff et al., 2018), providing evidence that the localizer task is ecologically valid. Thus, a breadth of evidence demonstrates that the *sentences > nonwords* contrast identifies a network that is engaged in language processing and appears to be a “natural kind” in the functional architecture of the human brain.

To identify MD regions, we used the *nonwords > sentences* contrast, targeting regions that increase their response with the more effortful reading of nonwords compared to that of sentences. This “cognitive effort” contrast robustly engages the MD network and can reliably localize it. Moreover, it generalizes across a wide array of stimuli and tasks, both linguistic and non-linguistic including, critically, contrasts targeting executive functions such as working-memory and inhibitory control (Fedorenko et al., 2013; Mineroff et al., 2018). **Supplementary Figures 1** and **2** demonstrate that the MD regions thus localized robustly respond to a difficulty (i.e. memory load) manipulation in a non-linguistic, spatial working-memory task (administered to a subset of participants in the current dataset).

*Main (story comprehension) task.* Participants listened to stories from the publicly available Natural Stories Corpus (Futrell et al., 2018). These stories were adapted from existing texts (fairy tales and short stories) to be “deceptively naturalistic”: they contained an over-representation of rare words and syntactic constructions embedded in otherwise natural linguistic context. Behavioral data indicate that these stories effectively manipulate predictive processing, as self-paced reading times from an independent sample show robust effects of surprisal (Futrell et al., 2018). Stories were recorded by two native English speakers (one male, one female) at a 44.1 kHz sampling rate, ranged in length from 4m46s to 6m29s (983-1099 words), and were played over scanner-safe headphones (Sensimetrics, Malden, MA).

### Data acquisition and preprocessing

#### Data acquisition

Structural and functional data were collected on a whole-body 3 Tesla Siemens Trio scanner with a 32-channel head coil at the Athinoula A. Martinos Imaging Center at the McGovern Institute for Brain Research at MIT. T1-weighted structural images were collected in 176 axial slices with 1mm isotropic voxels (repetition time (TR)=2,530ms; echo time (TE)=3.48ms). Functional, blood oxygenation level-dependent (BOLD) data were acquired using an EPI (echo-planar imaging) sequence with a 90° flip angle and using GRAPPA (GeneRalized Autocalibrating Partial Parallel Acquisition) with an acceleration factor of 2; the following parameters were used: thirty-one 4.4mm thick near-axial slices acquired in an interleaved order (with 10% distance factor), with an in-plane resolution of 2.1mm×2.1mm, FoV (field of view) in the phase encoding (Anterior>>Posterior) direction 200mm and matrix size 96mm×96mm, TR=2000ms and TE=30ms. The first 10s of each run were excluded to allow for steady state magnetization.

#### Spatial preprocessing

Data preprocessing was carried out with SPM5 and custom MATLAB scripts. Preprocessing of anatomical data included normalization into a common space (Montreal Neurological Institute (MNI) template), resampling into 2mm isotropic voxels, and segmentation into probabilistic maps of the gray matter, white matter (WM) and cerebrospinal fluid (CSF). Note that SPM was only used for preprocessing and basic first-level modeling, aspects that have not changed much in later versions; we used an older version of SPM because data for this study are used across other projects spanning many years and hundreds of participants, and we wanted to keep the SPM version the same across all the participants. Preprocessing of functional data included motion correction, normalization, resampling into 2mm isotropic voxels, smoothing with a 4mm FWHM Gaussian kernel and high-pass filtering at 200s.

#### Temporal preprocessing

Data from the story comprehension runs were additionally preprocessed using the CONN toolbox (Whitfield-Gabrieli and Nieto-Castañon, 2012) with default parameters, unless specified otherwise. Five temporal principal components of the BOLD signal time-courses from the WM were regressed out of each voxel’s time-course; signal originating in the CSF was similarly regressed out. Six principal components of the six motion parameters estimated during offline motion correction were also regressed out, as well as their first time derivative.

### Participant-specific functional localization of the language and MD networks

#### Modeling localizer data

A general linear model estimated the voxel-wise effect size of each condition in each experimental run of the localizer task. These effects were each modeled with a boxcar function (representing entire blocks/events) convolved with the canonical Hemodynamic Response Function (HRF). The model also included first-order temporal derivatives of these effects, as well as nuisance regressors representing entire experimental runs and offline-estimated motion parameters. The obtained beta weights were then used to compute the two functional contrasts of interest: *sentences > nonwords* for identifying language regions, and *nonwords > sentences* for identifying MD regions. These contrasts were computed only for voxels whose probability of belonging to the gray matter was greater than 1/3, based on the segmentation of the participant’s anatomical data. All other voxels were not considered further.

These group-level masks, in the form of binary maps, were used to constrain the selection of participant-specific fROIs. In particular, for each participant, 6 language fROIs were created by (i) intersecting each language mask with each individual participant’s unthresholded *t*-map for the *sentences > nonwords* contrast; and then (ii) choosing the 10% of voxels with highest *t*-scores in the intersection. Similarly, 20 MD fROIs were created by intersecting each MD mask with each participant’s unthresholded *t*-map for the *nonwords > sentences* contrast and selecting the 10% of voxels with the highest *t*-scores within each intersection. This top-10% criterion balances the trade-off between choosing only voxels that respond robustly to the relevant contrast and having a sufficient number of voxels in each fROI of each participant. Moreover, this criterion guarantees fROIs of identical size across participants (occupying 10% of each mask). Few exceptions to this criterion were made for those cases where less than 10% of the voxels in a mask showed a *t*-score greater than 0; here, we only included the subset of voxels with positive *t*-scores in the fROI, and excluded those voxels showing effects in the opposite direction.

**Figure 2:**
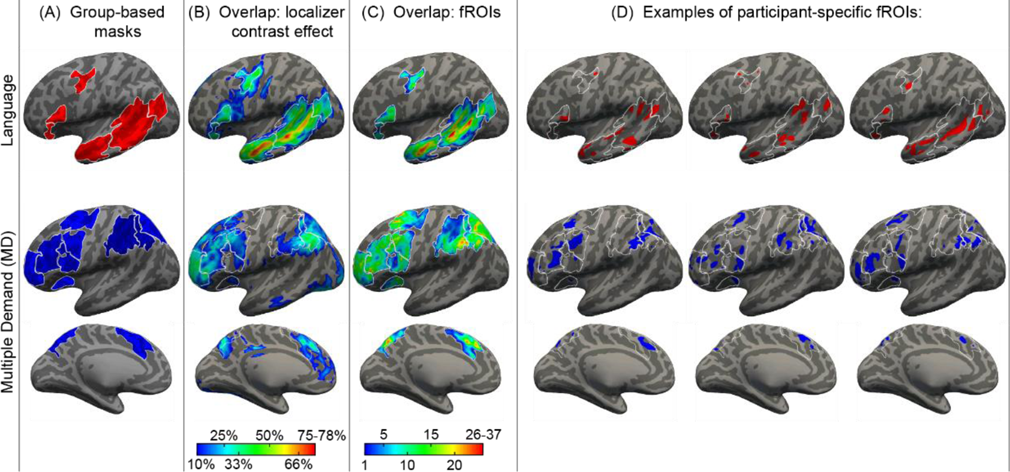
Defining participant-specific fROIs in the language (top) and MD (bottom) networks (only the left-hemisphere is shown). All images show approximated projections from functional volumes onto the surface of an inflated brain in common space. (A) Group-based masks used to constrain the location of fROIs. Contours of these masks are depicted in white on all brains in (B)-(D). (B) Overlap maps of localizer contrast effects (Sentence > Nonwords for the language network, Nonwords > Sentences for the MD network) across the 78 participants in the current sample (these maps were not used in the process of defining fROIs and are shown for illustration purposes). Each non gray-scale coordinate is colored according to the percentage of participants for whom that coordinate was among the top 10% of voxels showing the strongest localizer contrast effects across the nerocortical gray matter. (C) Overlap map of fROI locations. Each non gray-scale coordinate is colored according to the number of participants for whom that coordinate was included within their individual fROIs. (D) Example fROIs of three participants. Apparent overlap across language and MD fROIs within an individual is illusory and due to projection onto the cortical surface. Note that, because data were analyzed in volume (not surface) form, some parts of a given fROI that appear discontinuous in the figure (e.g., separated by a sulcus) are contiguous in volumetric space.

### Statistical analysis

#### Predictor definitions

To estimate word predictability in naturalistic data, we used an information-theoretic measure known as

*surprisal* (Shannon, 1948; Hale, 2001): the negative log probability of a word given its context. Surprisal can be computed in many ways, depending on the choice of probability model. Three previous naturalistic fMRI studies (Willems et al., 2015; Brennan et al., 2016; Lopopolo et al., 2017) searched for surface-level *n*-gram surprisal effects, using words and/or parts of speech as the token-level representation. In addition, two previous naturalistic fMRI studies (Brennan et al., 2016; Henderson et al., 2016) probed structure-sensitive PCFG surprisal measures (Hale, 2001; Roark et al., 2009). As discussed in the Introduction, results from these studies failed to converge on a clear answer as to the nature and functional location of surprisal effects. In this study, we used the following surprisal estimates:

- ***5-gram Surprisal*:** 5-gram surprisal for each word in the stimulus set from a KenLM (Heafield et al., 2013) language model with default smoothing parameters trained on the Gigaword 3 corpus (Graff et al., 2007). 5-gram surprisal quantifies the predictability of words as the negative log probability of a word given the four words preceding it in context.
- ***PCFG Surprisal*:** Lexicalized probabilistic context-free grammar surprisal computed using the incremental left-corner parser of van Schijndel et al. (2013) trained on a generalized categorial grammar (Nguyen et al., 2012) reannotation of Wall Street Journal sections 2 through 21 of the Penn Treebank (Marcus et al., 1993).

Models also included the control variables *Sound Power*, *Repetition Time (TR) Number*, *Rate*, *Frequency*, and *Network*, which were operationalized as follows:

- *Sound Power*: Frame-by-frame root mean squared energy (RMSE) of the audio stimuli computed using the Librosa software library (McFee et al., 2015).
- *TR Number*: Integer index of the current fMRI sample within the current scan.
- *Rate*: Deconvolutional intercept. A vector of ones time-aligned with the word onsets of the audio stimuli. *Rate* captures influences of stimulus *timing* independently of stimulus *properties* (see e.g., Brennan et al., 2016; Shain & Schuler, 2018).
- *Frequency*: Corpus frequency computed using a KenLM unigram model trained on Gigaword 3. For ease of comparison to surprisal, frequency is represented here on a surprisal scale (negative log probability), such that larger values index less frequent words (and thus greater expected processing cost).
- *Network*: Numeric predictor for network ID, 0 for MD and 1 for LANG.

Models additionally included the mixed-effects random grouping factors *Participant* and *fROI*. Prior to regression, all predictors were rescaled by their standard deviations in the training set except *Rate* (which has no variance) and *Network* (which is an indicator variable). Reported effect sizes are therefore in standard units.

#### Continuous-time deconvolutional regression

Naturalistic language stimuli pose a challenge for established statistical methods in fMRI because the stimuli (words) (1) are variably spaced in time and (2) do not temporally align with response samples recorded by the scanner. Previous approaches to address this issue have various drawbacks. Some fMRI studies of naturalistic language processing have assumed a canonical shape for the hemodynamic response function (Boynton et al., 1994) and used it to convolve stimulus properties into response-aligned measures (Willems et al., 2015; Brennan et al., 2016; Lopopolo et al., 2017). This approach is unable to account for regional variation in the shape of the hemodynamic response, even though the canonical HRF is known to be a poor fit to some brain regions (Handwerker et al., 2004). Discrete-time methods for data-driven HRF identification such as finite impulse response modeling (FIR; Dayal et al., 1996) and vector autoregression (VAR; Sims, 1980) are widely used to overcome the limitations of the canonical HRF for fMRI research (e.g., Friston et al., 1994; Harrison et al., 2003) but are of limited use in the naturalistic setting because they assume (multiples of) a fixed time interval between stimuli that does not apply to words in naturally-occurring speech. Some studies (e.g. Huth et al., 2016) address this problem by continuously interpolating word properties, resampling the interpolated signal so that it temporally aligns with the fMRI record, and fitting FIR models using the resampled design matrix. However, this approach can be distortionary in that word properties (e.g., surprisal) are not temporally continuous.

Our study employed a recently developed continuous-time deconvolutional regression (CDR) technique that accurately infers parametric continuous-time impulse response functions – such as the HRF – from arbitrary time series (Shain & Schuler, 2018, 2019). Because CDR is data-driven, it can address the potential impact of poor fit in the canonical HRF, and because it is defined in continuous time, it eliminates the need for distortionary preprocessing steps like continuous interpolation. CDR models in this study used the following two-parameter HRF kernel based on the widely-used double-gamma canonical HRF (Lindquist et al., 2009):

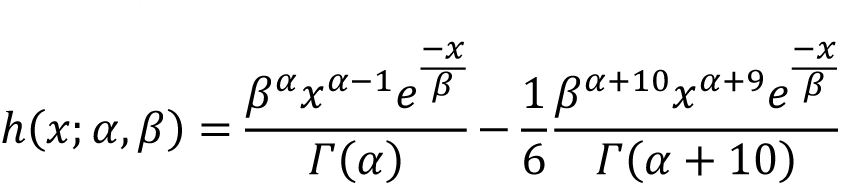

where α and β are initialized to the SPM defaults of 6 and 1, respectively. More complex kernels (e.g., that fit the amplitude of the second term, rather than fixing it at 1/6) were avoided because of their potential to overfit.

The parametric continuous-time nature of CDR is similar to that of models used, for example, by Kruggel & von Camon (1999), Kruggel et al. (2000), Miezin et al. (2000), Lindquist & Wager (2007), and Lindquist et al. (2009) for nonlinear estimation of gamma-shaped HRFs. The main advantages of CDR over these approaches are that (1) it exploits the Tensorflow (Abadi et al., 2015) and Edward (Tran et al., 2016) libraries for optimizing large-scale variational Bayesian computation graphs using state of the art estimation techniques from deep learning – this study used the Adam optimizer with Nesterov momentum (Kingma & Ba, 2014; Nesterov, 1983; Dozat, 2016); (2) it supports mixed effects modeling of effect coefficients and HRF parameters; and (3) it supports *parameter tying*, constraining the solution space by ensuring that all predictors share a common HRF shape in a given region (with potentially differing amplitudes). Predictors in these models were given their own coefficients (which rescale *h* above), but the parameters α and β of *h* were tied across predictors, modeling the assumption of a fixed-shape blood oxygenation response to neural activity in a given cortical region.

The CDR models applied in this study assumed improper uniform priors over all parameters in the variational posterior and were optimized using a learning rate of 0.001 and stochastic minibatches of size 1024. Following standard practice from linear mixed-effects regression (Bates et al., 2014), random effects were L2-regularized toward zero at a rate of 1.0. Convergence was declared when the loss was uncorrelated with training time by *t*-test at the 0.5 level for at least 250 of the past 500 training epochs.

For computational efficiency, predictor histories were truncated at 256 timesteps (words), which yields a maximum temporal coverage in our data of 48.34s (substantially longer than the effective influence of the canonical HRF). Prediction from the network used an exponential moving average of parameter iterates (Polyak, 1992) with a decay rate of 0.999, and models were evaluated using *maximum a posteriori* estimates obtained by setting all parameters in the variational posterior to their means. This approach is valid because all parameters are independent Gaussian in the CDR variational posterior (Shain & Schuler, 2018).

#### Model specification

The following CDR model specification was fitted to responses from each of the LANG and MD fROIs, where *italics* indicate predictors convolved using the fitted HRF and **bold** indicates predictors that were ablated for hypothesis tests:

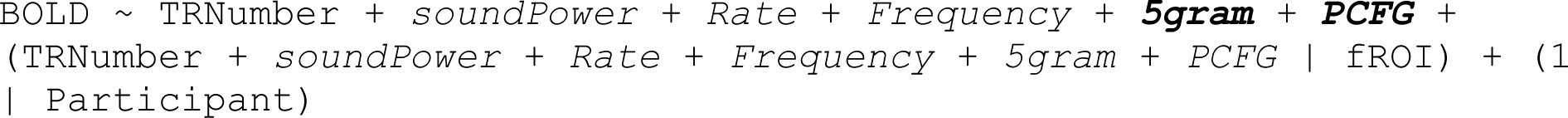

The random effect by fROI indicates that the model included zero-centered by-fROI random variation in response amplitude and HRF parameters for each functional region of interest. As shown, the model also included a random intercept by participant (the data do not appear to support richer by-participant random effects, e.g. including random slopes and HRF shapes, since such models explained no held-out variance in early analyses, indicating overfitting). The above model can test whether the surprisal variables help predict neural activation in a given cortical region. However, it cannot be used to compare the magnitudes of response to surprisal across networks (Nieuwenhuis et al., 2011). Therefore, we directly tested for a difference in influence by fitting the combined responses from both LANG and MD using the following model specification with the indicator variable *Network*:

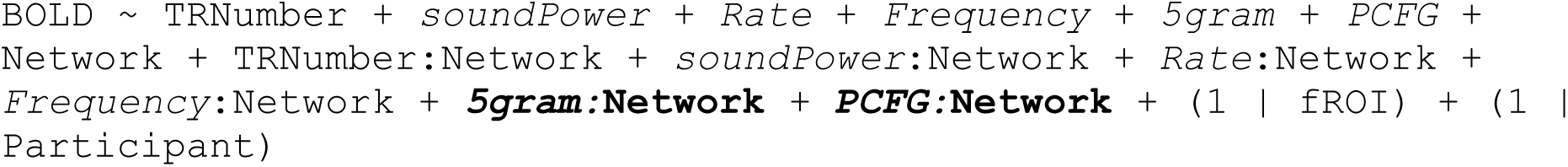

The random effects by fROI were simplified in comparison to that of the single-network models because the *Network* variable exactly partitions the fROIs. Thus ablated models can fully capture network differences as long as they have by-fROI random effects for surprisal. Indeed, initial tests showed virtually no difference in held-out likelihood between full and ablated combined models when those models included full by-fROI random effects despite large-magnitude estimates for the interactions with *Network* in the full model. Furthermore, the fitted parameters suggested that the by-fROI term was being appropriated in ablated models to capture between-network differences. In the full model, the *5-gram Surprisal* estimates for 50% of LANG fROI and 45% of MD fROI were positive, while in the model with *5gram:Network* ablated, 100% of LANG fROI and only 20% of MD fROI were positive, indicating that differences in response to *5-gram Surprisal* had been pushed into the by-fROI random term. For this reason, we used simpler models for the combined test, despite their insensitivity to by-fROI variation in HRF shape or response amplitude.

In interactions between *Network* and convolved predictors, the interaction was computed *following* convolution but prior to rescaling with that predictor’s coefficient. Thus, the interaction term represents the offset in the estimated coefficient from the MD network to the LANG network, as is the case for binary interaction terms in linear regression models.

Finally, exact deconvolution from continuous predictors like *Sound Power* is not possible, since such predictors do not have an analytical form that can be integrated. Instead, we sampled sound power at fixed intervals (100ms), in which case the event-based CDR procedure reduces to a Riemann sum approximation of the continuous convolution integral. Note that the word-aligned predictors (e.g. *5-gram Surprisal*) therefore have different timestamps than *Sound Power*, and as a result the history window spans different regions of time (up to 128 words into the past for the word-aligned predictors and up to 100ms × 128 = 12.8s of previous *Sound Power* samples).

#### Ablative statistical testing

In order to avoid confounds from (1) collinearity in the predictors and/or (2) overfitting to the training data, we followed a standard testing protocol from machine learning of evaluating differences in prediction performance on out-of-sample data using ablative non-parametric paired permutation tests for significance (Demšar, 2006). This approach can be used to assess the presence of an effect by comparing the prediction performance of a model that contains the effect against that of an ablated model that does not contain it. Specifically, given two pre-trained nested models, we computed the out-of-sample by-item likelihoods from each model over the evaluation set and constructed an empirical *p* value for the likelihood difference test statistic by randomly swapping by-item likelihoods *n* times (where *n*=10,000) and computing the proportion of obtained likelihood differences whose magnitude exceeded that observed between the two models. To ensure a single degree of freedom for each comparison, only fixed effects were ablated, with all random effects retained in all models.

The data partition was created by cycling TR numbers *e* into different bins of the partition with a different phase for each subject *u*:

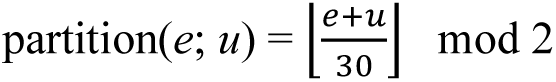

assigning output 0 to the training set and 1 to the evaluation set. Since TR duration is 2s, this procedure splits the BOLD times series into 60 second chunks, alternating assignment of chunks into training and evaluation sets with a different phase for each participant. Partitioning in this way allowed us to (1) obtain a model of each participant, (2) cover the entire time series, and (3) sub-sample different parts of the time series for each participant during training, while at the same time suppressing correlation between the training and evaluation responses by using a relatively long period of alternation (30 TRs or 60s).

### Accessibility

Access instructions for software and supplementary data needed to replicate these experiments (e.g. librosa, PyMVPA, CDR, KenLM, Gigaword 3, etc.) are given in the publications cited above. Post-processed fMRI timeseries are publicly available at the following URL: https://osf.io/eyp8q/. These experiments were not pre-registered.

### Results

The CDR-estimated mean double-gamma hemodynamic response functions (HRFs) for the LANG and MD networks are given in **Figure 3**, the estimated HRFs by fROI in LANG regions are shown in **Figure 4**, surprisal estimates and percent variance explained by region are given in **Tables 3** and **4**, and population-level effect estimates (i.e. areas under the estimated HRFs) are reported in **Table 5**. MD estimates by region are plotted in **Supplementary Figures 3 and 4**; they are of little relevance because they do not generalize (**Tables 4** & **6**). As shown, HRF shapes resemble but deviate slightly from the canonical HRF (Boynton et al., 1996) to varying degrees in each region, highlighting both consistency with HRF estimates established by prior research as well as the potential of CDR to discover subtle differences in HRF shape between cortical regions (Handwerker et al., 2004) in naturalistic data. The models find positive effects of similar strength for both *5-gram Surprisal* and *PCFG Surprisal* in LANG, and smaller effects of surprisal (even negative in the case of *5-gram Surprisal*) in MD.

**Figure 3:**
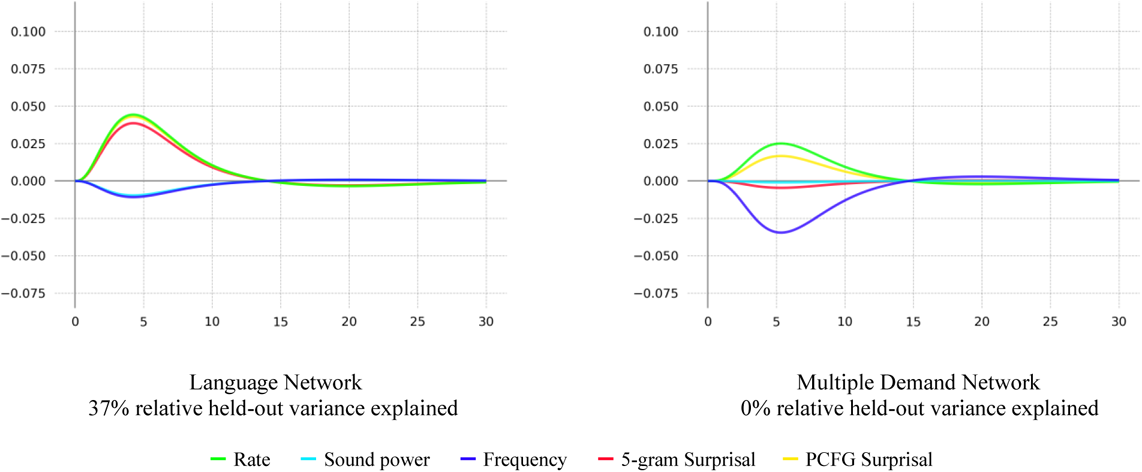
Estimated overall double-gamma hemodynamic response functions (HRFs) by network.

**Figure 4:**
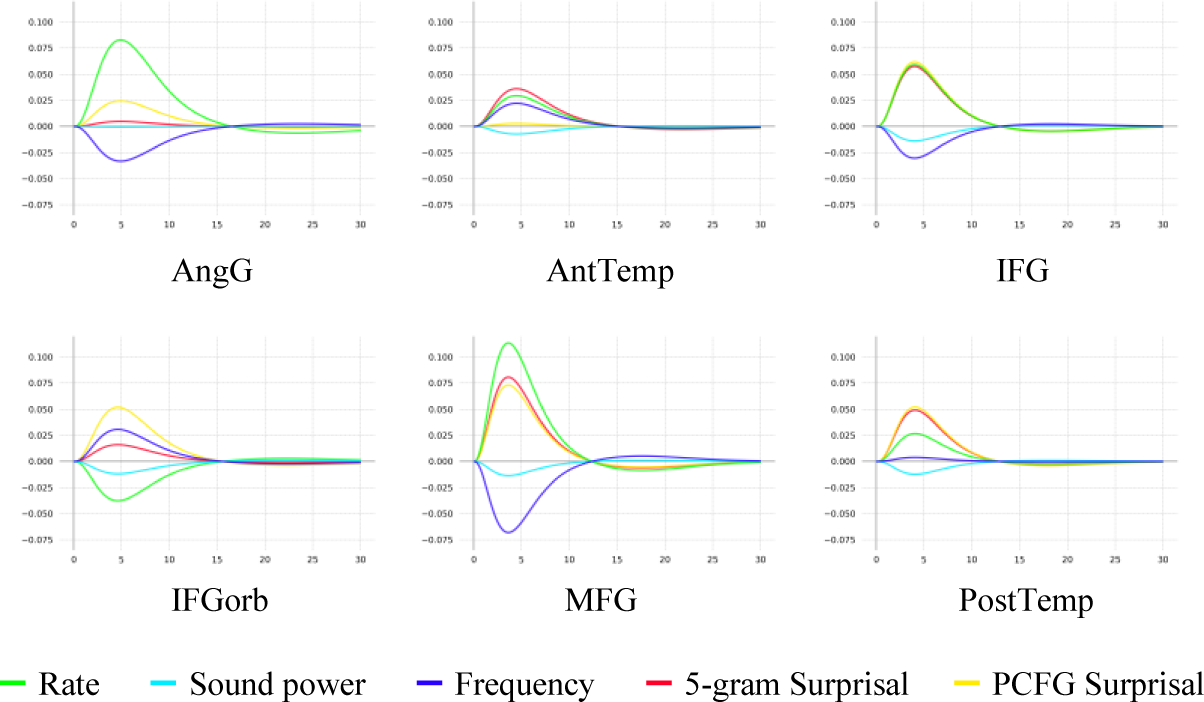
Estimated language-network HRFs by fROI.

**Table 3.** LANG surprisal estimates by fROI. Estimates given are the area under the fitted HRF. Models explain held-out variance in all regions but AngG.

| fROI | Hemisphere | 5-gram estimate | PCFG estimate | % Held-Out Variance Explained |
| --- | --- | --- | --- | --- |
| AngG | L | 0.030 | 0.156 | 0.0% |
| AntTemp | L | 0.215 | 0.017 | 5.1% |
| IFG | L | 0.287 | 0.309 | 2.2% |
| IFGorb | L | 0.010 | 0.318 | 1.3% |
| MFG | L | 0.382 | 0.346 | 2.3% |
| PostTemp | L | 0.242 | 0.258 | 6.1% |

**Table 4.** MD surprisal estimates by fROI. Estimates given are the area under the fitted HRF. Models explain no held-out variance in any region except left MFGorb.

| **fROI** | **Hemisphere** | **5-gram**<br>**estimate** | **PCFG**<br>**estimate** | **% Held-Out**<br>**Variance Explained** |
| --- | --- | --- | --- | --- |
| AntPar | L | 0.102 | -0.523 | 0.0% |
| IFGop | L | 0.009 | 0.141 | 0.0% |
| Insula | L | -0.200 | 0.284 | 0.0% |
| MFG | L | 0.074 | -0.026 | 0.0% |
| MFGorb | L | -0.215 | 0.252 | 0.5% |
| MidPar | L | 0.116 | -0.051 | 0.0% |
| mPFC | L | -0.125 | 0.257 | 0.0% |
| PostPar | L | 0.083 | -0.006 | 0.0% |
| PrecG | L | 0.078 | 0.048 | 0.0% |
| SFG | L | 0.180 | 0.025 | 0.0% |
| AntPar | R | 0.016 | -0.077 | 0.0% |
| IFGop | R | -0.011 | 0.075 | 0.0% |
| Insula | R | -0.185 | 0.227 | 0.0% |
| MFG | R | 0.058 | -0.006 | 0.0% |
| MFGorb | R | -0.004 | 0.019 | 0.0% |
| MidPar | R | 0.040 | -0.110 | 0.0% |
| mPFC | R | -0.321 | 0.440 | 0.0% |
| PostPar | R | -0.312 | 0.434 | 0.0% |
| PrecG | R | 0.034 | 0.118 | 0.0% |
| SFG | R | 0.066 | -0.034 | 0.0% |

**Table 5.** Model effect estimates.

|  | **Coefficient** |  |  |
| --- | --- | --- | --- |
| **Predictor** | **LANG** | **MD** | **Combined** |
| Sound Power | -0.055 | -0.006 | -0.003 |
| TR Number | -0.148 | 0.048 | -0.005 |
| Rate | 0.242 | 0.146 | 0.048 |
| Frequency | -0.060 | -0.199 | -0.134 |
| 5-gram Surprisal | 0.209 | -0.025 | 0.003 |
| PCFG Surprisal | 0.235 | 0.097 | 0.038 |
| Network | -- | -- | -1.32 |
| Sound Power by Network | -- | -- | -0.050 |
| TR Number by Network | -- | -- | -0.008 |
| Rate by Network | -- | -- | 0.269 |
| Frequency by Network | -- | -- | 0.040 |
| 5-gram Surprisal by Network | -- | -- | 0.212 |
| PCFG Surprisal by Network | -- | -- | 0.193 |

At the level of individual regions, the models explained held-out variance in all but one of the language fROIs (the exception was the AngG fROI). In contrast, the models explained no held-out variance in any but one MD fROI (the left MFGorb fROI). We leave these two exceptions to future research, but overall, the results demonstrate that surprisal effects are generally present throughout the language network and generally absent throughout the MD network. The differences between the individual-network models are largely replicated in the Combined model (**Table 5**), where main effects represent the estimated mean response in MD while interactions with *Network* represent the estimated difference in mean response between LANG and MD. As shown, Combined model estimates of both *5-gram:Network* and *PCFG:Network* are positive and large-magnitude, indicating that the model estimates these variables to yield greater increases in neural activity in LANG over MD.

**Table 6** reports model percent variance explained compared to a theoretical ceiling computed by regressing responses against responses from the same brain region in all other participants exposed to that stimulus. This ceiling is designed to quantify the variance that can be explained based on the stimuli alone, independently of inter-participant variation. As shown, models explain a substantial amount of the available variance in LANG. MD models explain no variance on the evaluation set, suggesting that the MD model did not learn generalizable patterns.

Because fROIs were modeled as random effects in these analyses, pairwise statistical testing of between-region differences in effect amplitude is not straightforward, and systematic investigation of regions / subnetworks within each broader functional network is left to future work. However, a qualitative examination of the by-region estimates suggests potentially interesting functional differences within the language network (**Table 3**). In particular, the IFG, MFG, and PostTemp fROIs all responded roughly equally to both measures of surprisal. The IFGorb fROI responded more to *PCFG* than *5-gram Surprisal* (an unexpected finding given that this is not the language region that is traditionally most strongly associated with syntactic processing; e.g., Friederici, 2011; Blank et al., 2016). The AngG fROI showed a similar pattern, but the models did not explain held-out variance for this fROI. And the AntTemp fROI responded more to *5-gram* than *PCFG Surprisal* (a finding which bears on debates about the functional role of this brain region in language processing, see Discussion). Although the differences in effect sizes between the two surprisals are significant in each of IFGorb, AngG, and AntTemp by Monte Carlo estimated credible intervals tests, such tests are anticonservative in CDR (Shain & Schuler, 2019).

**Tables 7-9** show the main finding of this study: fixed effects for *5-gram Surprisal* and *PCFG Surprisal* significantly improve held-out likelihood in the LANG network over a model containing neither, as well as over one another. The difference in effect size between the LANG and MD networks is statistically significant, as shown by the significant likelihood improvements yielded by interactions of the surprisal variables with *Network*.

**Table 6.**
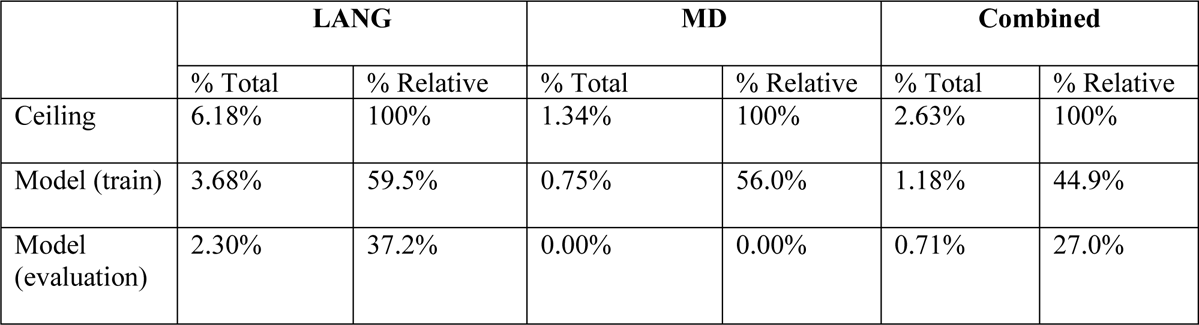
Model percent variance explained compared to a “ceiling” linear model regressing against the mean response of all other participants for a particular story/fROI. “% Total” columns show absolute percent variance explained, while “% Relative” columns show percent variance explained relative to the ceiling.

|  | **LANG** |  | **MD** |  | **Combined** |  |
| --- | --- | --- | --- | --- | --- | --- |
|  | % Total | % Relative | % Total | % Relative | % Total | % Relative |
| Ceiling | 6.18% | 100% | 1.34% | 100% | 2.63% | 100% |
| Model (train) | 3.68% | 59.5% | 0.75% | 56.0% | 1.18% | 44.9% |
| Model (evaluation) | 2.30% | 37.2% | 0.00% | 0.00% | 0.71% | 27.0% |

**Table 7.** LANG result. Significance in LANG by paired permutation test of log-likelihood improvement on the evaluation set from including a fixed effect for each of *5-gram Surprisal* and *PCFG Surprisal*, over (1) a baseline with neither fixed effect and (2) baselines containing the other fixed effect only. The *Effect Estimate* column shows the estimated effect size from the model containing the fixed effect (i.e. the area under the estimated HRF).

| Comparison | $p$ | LL Improvement | Effect Estimate |
| --- | --- | --- | --- |
| 5-gram over neither | 0.0001*** | 182 | 0.307 |
| PCFG over neither | 0.0001*** | 183 | 0.352 |
| 5-gram over PCFG | 0.0001*** | 61 | 0.209 |
| PCFG over 5-gram | 0.0001*** | 61 | 0.235 |

**Table 8.** MD result. Significance in MD by paired permutation test of log-likelihood improvement on the evaluation set from including a fixed effect for each of *5-gram Surprisal* and *PCFG Surprisal*, over (1) a baseline with neither fixed effect and (2) baselines containing the other fixed effect only. A *p*-value of 1.0 is assigned by default to comparisons in which held-out likelihood improved under ablation. The *Effect Estimate* column shows the estimated effect size from the model containing the fixed effect (i.e. the area under the estimated HRF).

| Comparison | $p$ | LL Improvement | Effect Estimate |
| --- | --- | --- | --- |
| 5-gram over neither | 0.137 | 3 | 0.019 |
| PCFG over neither | 1.0 | -29 | 0.081 |
| 5-gram over PCFG | 1.0 | -8 | -0.025 |
| PCFG over 5-gram | 1.0 | -40 | 0.097 |

**Table 9.** Combined result. Significance in the combined data by paired permutation test of log-likelihood improvement on the evaluation set from including a fixed interaction for each of *5-gram Surprisal* and *PCFG Surprisal* with *Network*, over (1) a baseline with neither fixed interaction and (2) baselines containing the other fixed interaction only. The *Effect Estimate* column shows the estimated interaction size from the model containing the fixed interaction (i.e. the difference in effect estimate between LANG and MD).

| Comparison | $p$ | LL Improvement | Effect Estimate |
| --- | --- | --- | --- |
| 5-gram:Network over neither | 0.0001*** | 144 | 0.212 |
| PCFG:Network over neither | 0.0001*** | 144 | 0.193 |
| 5-gram:Network over PCFG:Network | 0.0001*** | 53 | 0.301 |
| PCFG:Network over 5-gram:Network | 0.0001*** | 53 | 0.317 |

As shown in **Figure 3**, the effects signs for *Frequency* in both networks are negative. The lack of a positive effect of *Frequency* is not what would be expected if word frequency modulated neural activity (Staub, 2015), but it is consistent with recent naturalistic behavioral evidence against distinct effects of frequency and predictability (Shain, 2019), as well as with previous theoretical claims that apparent frequency effects are underlyingly effects of predictability (Levy, 2008). Negative effects like these indicate suppression of the BOLD response and pose a challenge for interpretation (Harel et al., 2002). Prior work has suggested that such negative effects can arise from increased processing load elsewhere in the brain through hemodynamic factors (“vascular steal”) (Lee et al., 1995; Saad et al., 2001; Harel et al., 2002; Kannurpattie et al., 2004) and/or neuronal ones such as inhibition by an attention mechanism (Smith et al., 2000; Shmuel et al., 2002; Shmuel et al., 2006). The means by which such mechanisms might give rise to negative frequency effects in these experiments are not currently clear. Since frequency effects are not central to our present research question, we leave targeted investigation of their existence and direction to future research.

**Figure 7** and **Table 10** assess the generalizability of surprisal effects across participants. **Figure 7** shows most by-participant improvements clustered around a positive median, without strong visual indication of large-magnitude positive outliers that might exclusively drive the effect. This intuition is quantified in **Table 10**. As shown, held-out likelihood improves for most participants in all comparisons. Furthermore, at least 5 of the most responsive participants in each comparison can be removed without changing the significance of the effect. Participant removal is a stringent criterion not only because it excludes the most responsive participants from consideration but also because it reduces the power of the permutation test by shrinking the evaluation set. These participant-level analyses demonstrate that surprisal effects in LANG are not merely driven by a small number of outlier participants.

**Figure 7:**
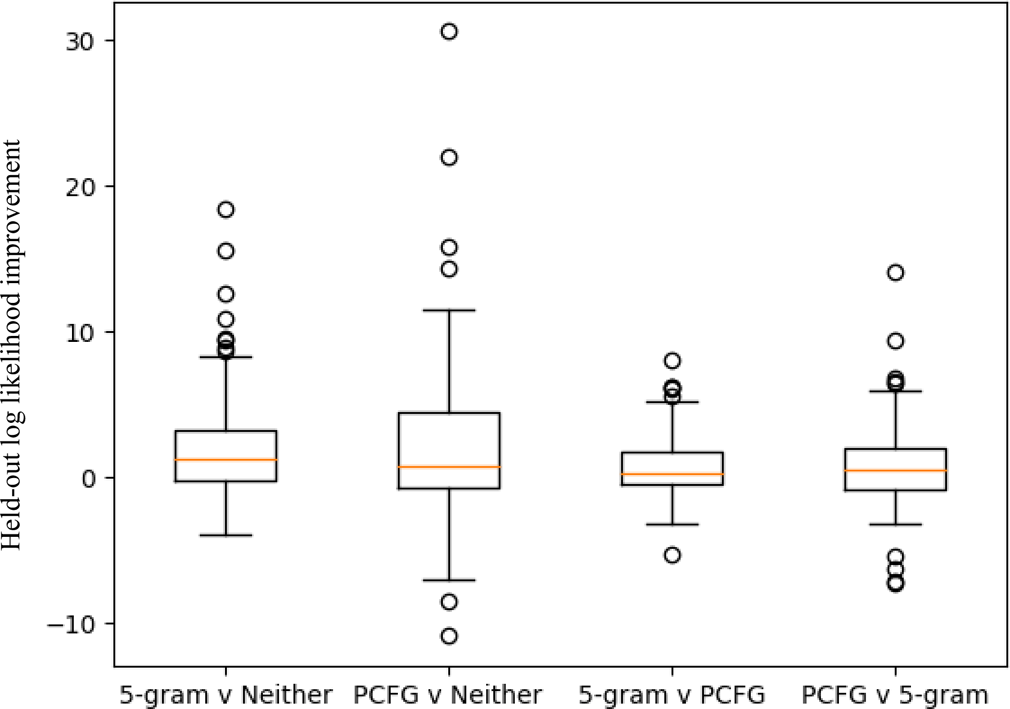
LANG likelihood improvement by participant. Spread of by-participant likelihood improvements in each comparison. Most improvements are positive, and effects are not driven by large positive outliers (see Table 10).

**Table 10.** Generality of LANG surprisal effects across participants. Median likelihood improvement in LANG on the evaluation set by participant, percent of participants whose held-out predictions improved due to surprisal effects, and the number of participants with the largest held-out improvement whose data can be removed without changing the significance of the effect at a 0.05 level. Held-out likelihood improves for most participants in every comparison, and at least 5 of the most responsive participants can be removed in each comparison without changing the significance of the effect.

| Comparison | Median LL Improvement by Participant | % Participants Improved | Num Removable Participants |
| --- | --- | --- | --- |
| 5-gram over neither | 1.236 | 71.8% | 19 |
| PCFG over neither | 0.732 | 64.1% | 14 |
| 5-gram over PCFG | 0.335 | 61.5% | 7 |
| PCFG over 5-gram | 0.498 | 60.3% | 5 |

### Discussion

The current study examined signatures of predictive processing during naturalistic story comprehension in two functionally distinct cortical networks: the domain-specific language (LANG) network, and the domain-general multiple demand (MD) network. Specifically, we tested which of these networks increased their responses with lower word predictability, operationalized using both 5-gram and probabilistic context-free grammar (PCFG) surprisal. The main results, yielded by continuous-time deconvolutional regression (CDR) analysis of surprisal effects in the two networks, are shown in **Tables 7-9**: in LANG, both *5-gram Surprisal* and *PCFG Surprisal* have positive effects that yield statistically significant improvements to held-out likelihood, both over a baseline containing neither fixed effect as well as over one another. By contrast, in MD, neither surprisal effect is significant in any comparison. A direct test for a difference in surprisal effects across the two networks (**Table 9**) shows that the interactions of both surprisals with network are positive and statistically significant, indicating that the BOLD response to both surface-level (5-gram) and structural (PCFG) word predictability is larger in LANG than MD. These results are over a baseline that includes an effect for lexical frequency (log unigram probability), which is notable given the strong natural correlation between surprisal and frequency, both generally (Demberg & Keller, 2008) and in the current experimental materials (*r* = 0.78 overall). This finding suggests that the surprisal effects reported here are indeed driven by predictive coding and not merely by the cost of retrieving infrequent words. Together, these results demonstrate that predictive coding for upcoming words is primarily a canonical computation carried out by domain-specific cortical circuits, rather than by feedback from higher, domain-general executive control circuits, and that these predictions depend on both surface-level and structural information sources. Our finding of a generalized effect of *PCFG Surprisal* throughout the language network aligns with prior findings of evidence for linguistic prediction (e.g. Kuperberg et al., 2000; Baumgaertner et al., 2002; Friederici et al., 2003; Obleser et al., 2007) and syntactic processing (e.g., Blank et al., 2016; see Zaccarella et al., 2017 for review) in these regions, but suggests that prior evidence of linguistic prediction effects in MD (e.g. Kuperberg et al., 2000; Baumgaertner et al., 2002; Gold et al., 2006; Bonhage et al., 2015; Hartwigsen et al., 2017) may have been influenced by the use of artificially constructed linguistic stimuli and/or task artifacts.

The aforementioned debate about the compartmentalization of language processing has largely focused on controlled experimental paradigms, which are prone to induce task artifacts that confound functional differentiation of neural structures. By showing strong prediction-based functional differentiation between the LANG and MD networks during naturalistic language comprehension, the present study provides evidence that predictive coding for language is primarily carried out by language-specific rather than domain-general mechanisms.

The finding that surprisal computed by marginalizing over syntactic structures (*PCFG Surprisal*) modulates the LANG response independently of surface-level *n*-gram surprisal is evidence that participants are indeed computing such structures during incremental sentence processing (Hale, 2001; Levy, 2008; Fossum & Levy, 2012; Rasmussen & Schuler, 2018) and is inconsistent with previous arguments that the human sentence processing response is largely insensitive to such structures (Frank & Bod, 2011; Frank et al., 2012; Frank & Christiansen, 2018). At the same time, the finding that *5-gram Surprisal* modulates the LANG response independently of *PCFG Surprisal* is evidence that the human sentence processing mechanism is sensitive to word co-occurrence patterns in ways that are not well captured by a strictly context-free parser. This suggests either (1) that the human parser is not strictly context-free (see e.g., tree-adjoining grammars, Joshi, 1985; combinatory categorial grammars, Steedman, 2000; and other context-sensitive grammar formalisms for natural language), or (2) that participants track both hierarchical structure and word co-occurrence patterns separately and simultaneously when generating predictions, and that these two kinds of processes take place in overlapping brain areas.

Evaluating these hypotheses is left to future work. The lack of structured prediction effects in MD is of interest given prior proposals that ground structural effects in constraints on working memory (Abney & Johnson, 1991; Resnik, 1992; Rasmussen & Schuler, 2018). These theories view the processing of hierarchical language structures as a special case of a domain general capacity for hierarchic sequential prediction (Botvinick, 2007), which is at least consistent with the hypothesis that the resources recruited for prediction are also domain general (see e.g. Smith & Levy, 2013). However, to the extent that the memory resources used for prediction are also expected to activate in response to prediction error (e.g., by undergoing model revision, Chao et al., 2018, see Introduction), the failure to find such a signal in MD suggests that these memory resources may also be specific to the functional language network, rather than domain general (e.g., Caplan & Waters, 1999; Matchin, 2017).

Estimates at the fROI level shed light on results from prior naturalistic fMRI experiments (Willems et al., 2015; Brennan et al., 2016; Henderson et al., 2016; Lopopolo et al., 2017). We found strong effects of both surface-level and structural estimates of word predictability in roughly the union of left-hemisphere language regions for which such effects have been reported in prior work (e.g., temporal and inferior frontal regions). At the same time, we did not find clear evidence of predictive coding in regions linked with the multiple demand network, like superior frontal gyrus (cf. Lopopolo et al., 2017), in part because our use of held-out significance tests helped us avoid reporting MD surprisal effects that fail to generalize (e.g., left-hemisphere SFG, **Table 4**). The lack of held-out testing in earlier studies may therefore have contributed to prior findings of surprisal effects in MD regions. Finally, we obtained significant positive effects for surprisal implementations in language regions that have previously been reported null or negative (e.g., lexicalized trigrams in IFG and posterior temporal cortex or PCFG surprisal in IFG, per Brennan et al., 2016; PCFG surprisal in the temporal lobe, per Henderson et al., 2016). It is possible that the size of the present study increased sensitivity to these effects, since studies using less data are more likely to yield sign and magnitude errors (Gelman & Carlin, 2014). The picture that emerges more clearly from our results than from those of prior studies is of a predictive coding mechanism that is specific to the functional language network, generalized throughout it, and sensitive to both surface-level word co-occurrence patterns and hierarchical structure.

Our emphasis on structural influences on *prediction*, rather than sensitivity to syntactic structure more generally, is a possible explanation for one apparent discrepancy between our results and those of some previous studies. In particular, we do not find evidence of *PCFG Surprisal* effects in the AntTemp language fROI (although the *PCFG Surprisal* estimate is positive, the model explains no held-out variance in AntTemp), whereas numerous previous studies have argued for syntactic effects in left anterior temporal cortex, both using hand-constructed stimuli (Mazoyer et al., 1993; Stowe et al., 1998; Friederici et al., 2000; Vandenberghe et al., 2002; Dronkers et al., 2004; Humphries et al., 2006; Rogalsky & Hickok, 2009; Pallier et al., 2011; Brennan & Pylkkänen, 2012; Nelson et al., 2017) and naturalistic stimuli (Brennan et al., 2010; Brennan & Pylkkänen, 2017; Bhattasali et al., 2018, 2019). The role of left anterior temporal cortex in syntactic processing has been called into question by an absence of syntactic deficits in patients with anterior temporal damage (e.g., Wilson et al., 2012), and some have argued that parts of the anterior temporal lobe primarily carry out lexical and semantic processing, including perhaps semantic composition (e.g., Bemis & Pylkkänen, 2011), rather than syntactic structure building (Visser et al., 2010; Wilson et al., 2014; Lambon Ralph et al., 2017; see also Matchin et al., 2018). However, even granting that left anterior temporal cortex is implicated in syntactic processing, prior studies by and large have focused on structural measures that are arguably integrative in nature (syntactic node count, number of parser operations, etc.) or have used manipulations that are too broad to target prediction vs. integration (sentences vs. list of words or “Jabberwocky” sentences). Indeed, claims about syntactic processing in left anterior temporal cortex tend to focus on composition rather than on structured prediction. Our results thus do not preclude a role for left anterior temporal cortex in structure-building broadly construed; they simply fail to show strong evidence in this brain area of effects of structural context on word predictability. Prior studies of structured prediction effects in left anterior temporal cortex have yielded mixed results; although Brennan et al. (2016) found evidence of part-of-speech *n*-gram and PCFG surprisal in anterior temporal cortex over bi- and tri-gram effects, Lopopolo et al. (2017) did not find a response to part-of-speech *n*-gram surprisal, and the response to syntactic PCFG surprisal in Henderson et al. (2016) was too weak to achieve significance. Prediction effects based on *lexical* context in left anterior temporal cortex (i.e. lexical *n*-grams) are better attested (Willems et al., 2015; Lopopolo et al., 2017), and some have explicitly argued that left anterior temporal cortex plays a central role in lexical-semantic prediction (Lau et al., 2016). Our findings in the AntTemp fROI (large effects of *5-gram Surprisal* in the AntTemp language fROI) contribute to this debate, suggesting that lexical prediction does occur in left anterior temporal cortex (among other regions) while syntactic prediction likely occurs elsewhere. Left anterior temporal cortex may therefore be an important object of study in teasing apart predictive vs. integrative processing during language comprehension, and further investigation is warranted.

In summary, our findings based on a large-scale naturalistic fMRI experiment support a view of linguistic prediction as implemented by domain-specific cortical circuits, sensitive to both surface-level and syntactic information sources, and generalized across the functional language network.

## Acknowledgments

This research was supported by NIH award R00-HD-057522 (E.F.) and by National Science Foundation grant #1816891 (W.S.). All views expressed are those of the authors and do not necessarily reflect the views of the National Science Foundation. E.F. was additionally supported by NIH awards R01-DC-016607 and R01-DC-016950 and by a grant from the Simons Foundation via the Simons Center for the Social Brain at MIT. The authors would also like to acknowledge the Athinoula A. Martinos Imaging Center at the McGovern Institute for Brain Research at the Massachusetts Institute of Technology (MIT), and the support team (especially Steven Shannon and Atsushi Takahashi). The authors also thank Nancy Kanwisher and Ted Gibson for recording the stories, and EvLab members for help with data collection, especially Zach Mineroff and Alex Paunov.

