## Supplemental Figures for "fMRI reveals language-specific predictive coding during naturalistic sentence comprehension" (from another version of this preprint: version 1, 2019-07-28)

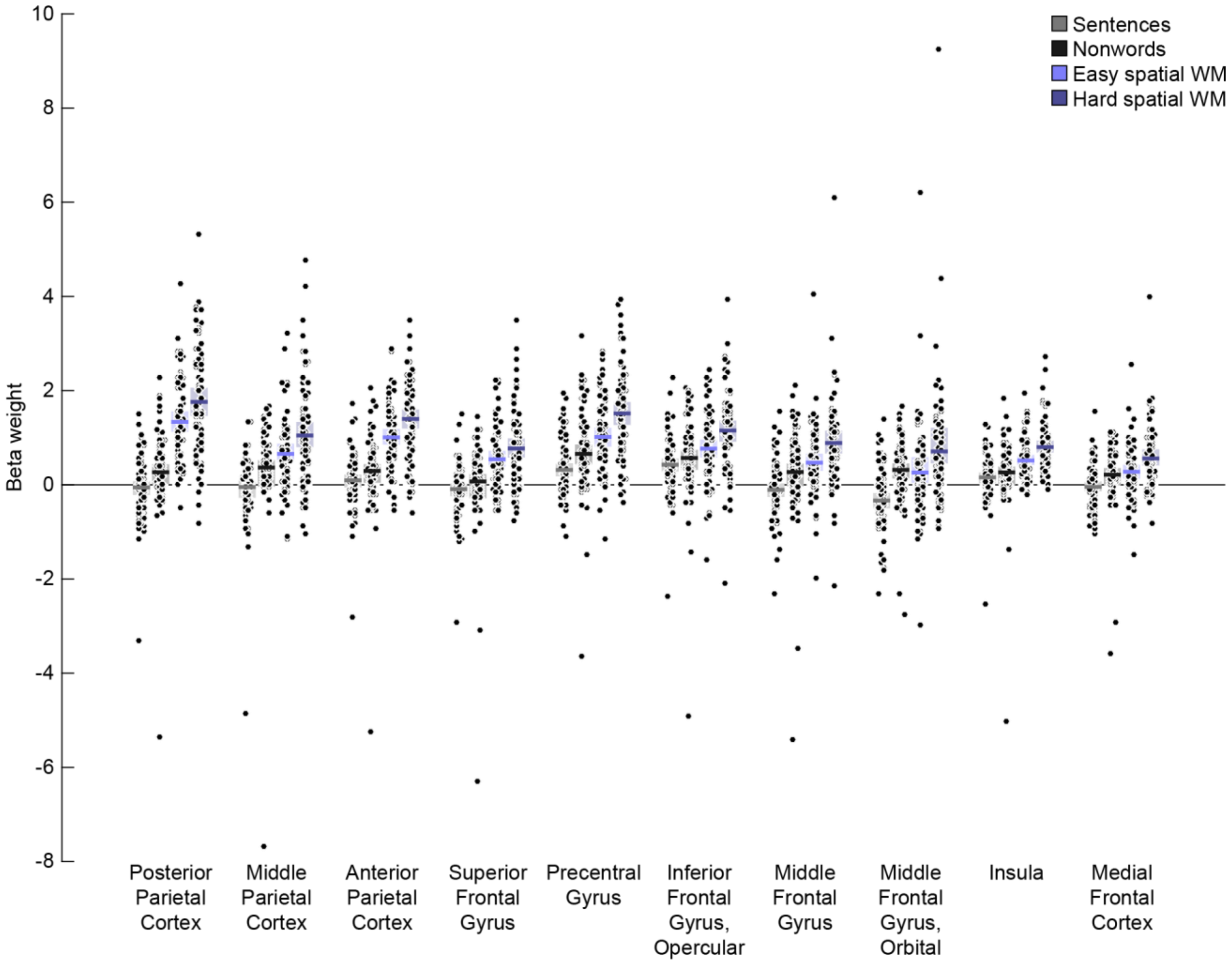


Supplementary Figure 1: Response profiles of participant-specific fROIs in the left-hemispheric MD network. Response magnitudes are shown (i) for the conditions of the localizer task, i.e., sentences (gray) and nonwords (dark gray) (estimated using across-runs cross-validation); and (ii) for a spatial, working-memory task that includes an easier (light blue) and a harder (dark blue) condition. All fROIs show the signature response to the difficulty manipulation, i.e., a significant Hard > Easy effect for the spatial WM task ($p < 10^{-4}$ following FDR correction). Dots represent data from individual participants, thick lines represent means, and rectangles surrounding the means represent 95% bootstrap confidence intervals.

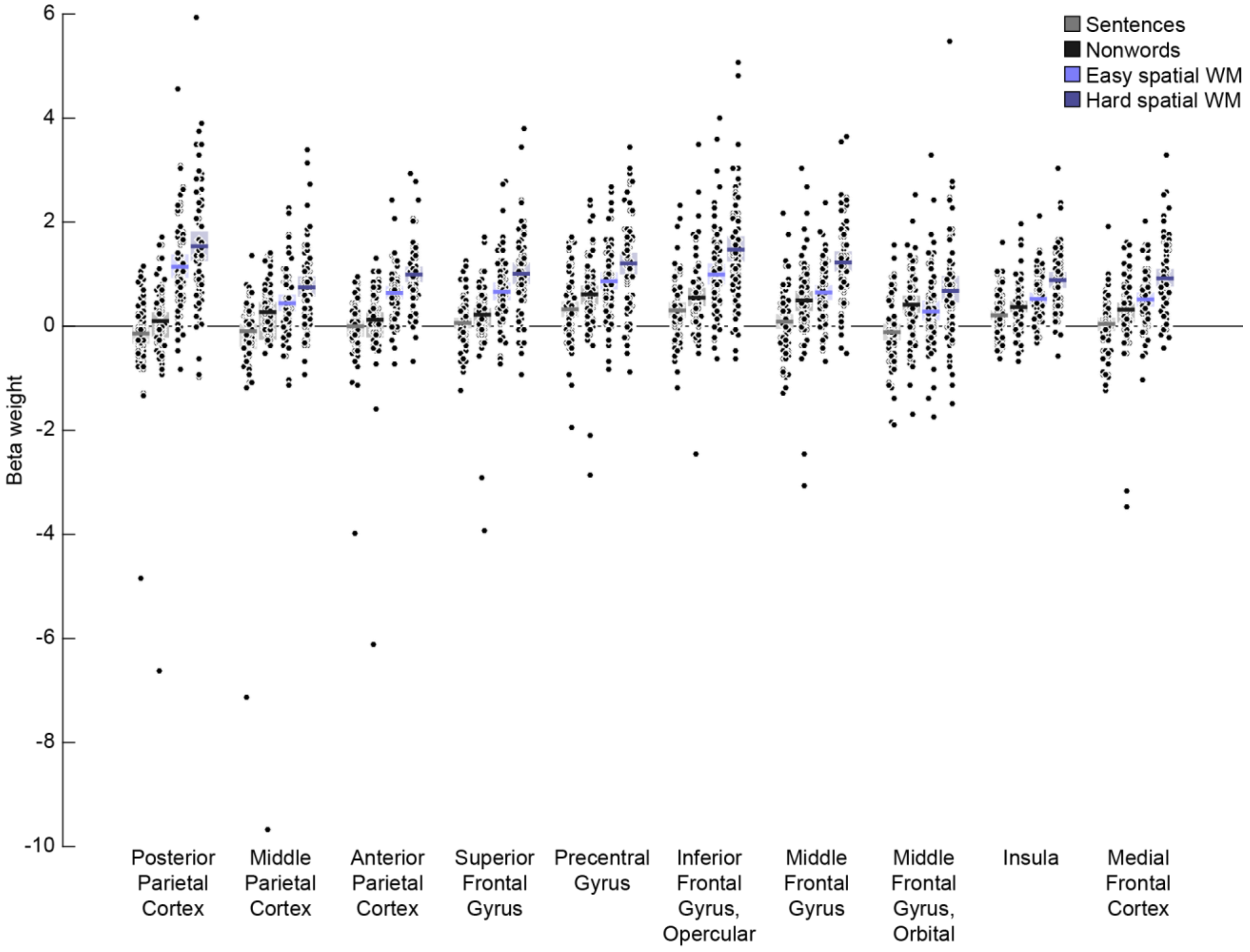


Supplementary Figure 2: Response profiles of participant-specific fROIs in the right-hemispheric MD network. Conventions are the same as in Supplementary Figure 1.

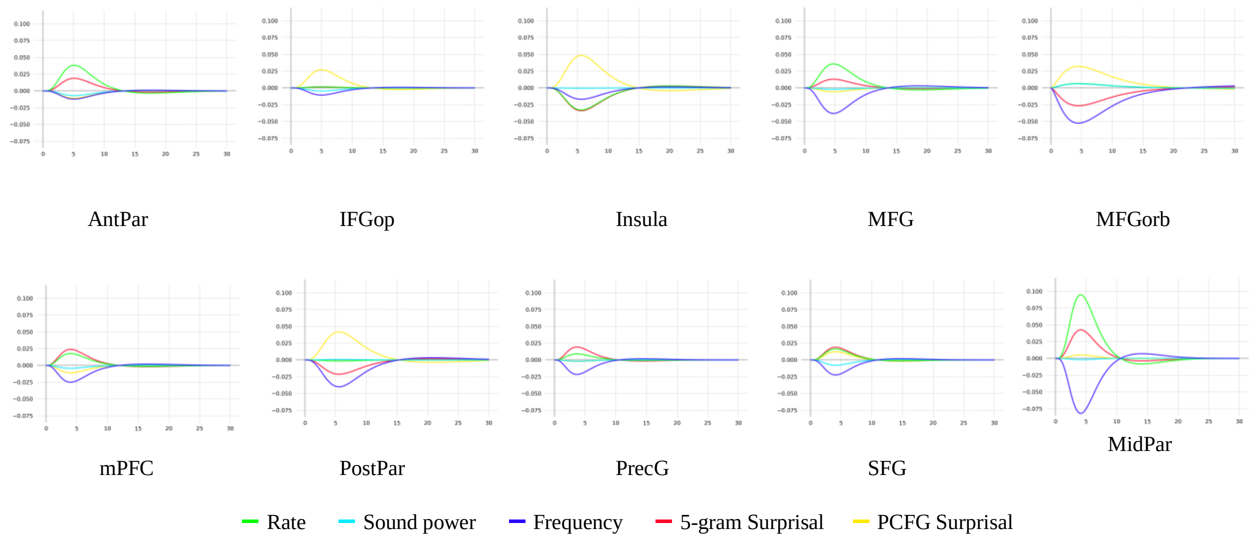


Supplementary Figure 3: Estimated left-hemisphere MD network HRFs by fROI

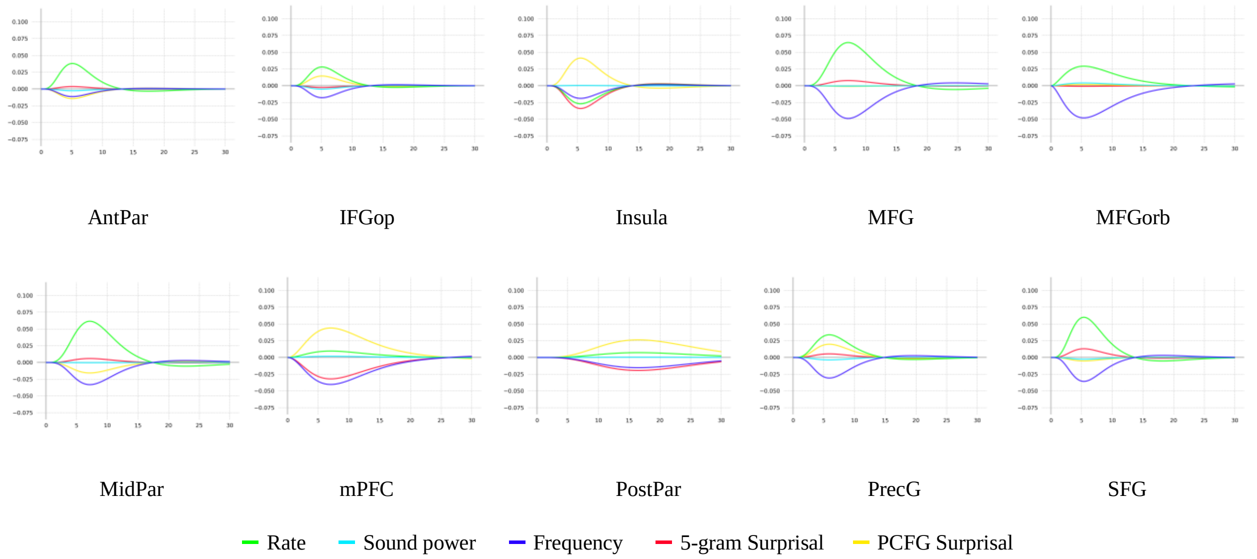


Supplementary Figure 4: Estimated right-hemisphere MD network HRFs by fROI
